# Inferring social structure and its drivers from refuge use in the desert tortoise, a relatively solitary species

**DOI:** 10.1101/025494

**Authors:** Pratha Sah, Kenneth E. Nussear, Todd C. Esque, Christina M. Aiello, Peter J. Hudson, Shweta Bansal

## Abstract

For several species, refuges (such as burrows, dens, roosts, nests) are an essential resource for protection from predators and extreme environmental conditions. Refuges also serve as focal sites for social interactions including mating, courtship and aggression. Knowledge of refuge use patterns can therefore provide information about social structure, mating and foraging success, as well as the robustness and health of wildlife populations, especially for species considered to be relatively solitary. In this study, we construct networks of burrow use to infer social associations in a threatened wildlife species typically considered solitary - the desert tortoise. We show that tortoise social networks are significantly different than null networks of random associations, and have moderate spatial constraints. We next use statistical models to identify major mechanisms behind individual-level variation in tortoise burrow use, popularity of burrows in desert tortoise habitat and test for stressor-driven changes in refuge use patterns. We show that seasonal variation has a strong impact on tortoise burrow switching behavior. On the other hand, burrow age and topographical condition influence the number of tortoises visiting a burrow in desert tortoise habitat. Of three major population stressors affecting this species (translocation, drought, disease), translocation alters tortoise burrow switching behavior, with translocated animals visiting fewer unique burrows than residents. In a species that is not social, our study highlights the importance of leveraging refuge use behavior to study the presence of and mechanisms behind non-random social structure and individual-level variation. Our analysis of the impact of stressors on refuge-based social structure further emphasizes the potential of this method to detect environmental or anthropogenic disturbances.

Significance statement: Adaptive and social behavior that affects fitness is now being increasingly incorporated in the conservation and management of wildlife species. However, direct observations of social interactions in species considered to be solitary are difficult, and therefore integration of behavior in conservation and management decisions in such species has been infrequent. For such species, we propose quantifying refuge use behavior as it can provide insights towards their (hidden) social structure, establish relevant contact patterns of infectious disease spread, and provide early warning signals of population stressors. Our study highlights this approach in a long-lived and threatened species, the desert tortoise. We provide evidence towards the presence of and identify mechanisms behind the social structure in desert tortoises formed by their burrow use preferences. We also show how individuals burrow use behavior responds to the presence of population stressors.

## Introduction

Social structure of wildlife populations is typically derived from observational studies on direct social interactions [e.g. affiliative interactions in primates (Griffin and Nunn 2011; MacIntosh et al. 2012), group association in dolphins (Lusseau et al. 2006) and ungulates (Cross et al. 2004; Vander Wal et al. 2012), food sharing in vampire bats (Carter and Wilkinson 2013)]. In relatively solitary species, individuals spend a considerable amount of time alone and have minimal direct interactions with conspecifics except during mating and occasional aggressive encounters (Scott and Carrington 2011). Examples of such species include raccoons, red foxes, orangutans, and some species of bees, wasps and bats. For these wildlife populations, social interactions may be limited to certain areas within their habitat, such as refuges (e.g., roost, den, burrow, nest) or watering holes that provide increased opportunities of direct contact between individuals. Monitoring these resources can therefore help establish relevant social patterns among individuals.

In addition to establishing social structure, refuges provide shelter, protection from predators and serve as sites for nesting and mating. Refuge use patterns of individuals are therefore central to survival, mating and foraging success and can serve as efficient indicators of population disturbances. Unlike traditional population dynamics indicators such as mortality and birth rate, refuge use behavior can respond instantaneously to sub-optimal conditions (Morris et al. 2009; Berger-Tal et al. 2011). Altered patterns of refuge use may thus indicate a disturbance or change in population fitness and provide an early warning to conservation biologists. Changes in habitat or refuge use have indeed been linked to the presence of natural population stressors such as increased predation (van Gils et al. 2009), drought (Kerr and Bull 2006; Gough et al. 2012) and disease transmission risk (Behringer and Butler IV 2010), as well as anthopogenic population stressors of translocation (Jachowski et al. 2012) and urbanization (Moule et al. 2015).

While the importance of refuge use in social interactions, survival and mating success, as well as indicators of environmental and anthropogenic stressors has been long appreciated, biologists are only beginning to understand individual level heterogeneity in refuge use and its population-level consequences in relatively solitary species (Fortuna et al. 2009; Leu et al. 2010; Godfrey 2013). The general absence of studies quantifying pairwise interactions due to preferences in refuge-use implies a lack of knowledge of the baseline social organization that could be used to evaluate changes in robustness or health of these wildlife populations. To overcome these shortcomings we explore a modeling framework that combines network theory with statistical models to infer the presence of and mechanisms behind the social organization in the desert tortoise, *Gopherus agassizii*, formed by their refuge-use preferences. The desert tortoise is a long-lived, terrestrial species that occurs throughout the Mojave Desert north and west of the Colorado River. Individuals of this species use subterranean burrows as an essential adaptation to obtain protection from temperature extremes and predators. Because tortoises spend a majority of their time either in or near burrows, most of their social interactions are associated with burrows (Bulova 1994).

Social behavior in desert tortoises is not well understood, though evidence suggests the presence of dominance hierarchies (Niblick et al. 1994; Bulova 1997) which may influence social structure and burrow choice in desert tortoises. In addition to social hierarchies, previous research suggests factors such as sex (Harless et al. 2009), age (Wilson et al. 1999), season (Bulova 1994); and environmental conditions (Duda et al. 1999; Franks et al. 2011) may influence burrow use in desert tortoises. If conspecific cues and environmental factors exhibit strong influence on burrow use, population stressors impacting these characteristics could alter typical burrow behavior. The two major population threats that have been identified in desert tortoise populations include upper respiratory tract disease (URTD) caused by *Mycoplasma agassizii* and *Mycoplasma testudineum* (Brown et al. 1994; Sandmeier et al. 2009; Jacobson et al. 2014), and extreme environmental conditions, particularly drought (Longshore et al. 2003; Lovich et al. 2014). In addition to these threats, the primary management strategy in desert tortoises is to translocate animals out of areas affected by anthropogenic disturbances (Department of the Interior 2011). Translocation in other reptilian species, however, has had limited success due to high rates of mortality (Dodd and Seigel 1991; Germano and Bishop 2009) and may also act as a population stressor. In desert tortoises, all three population stressors have been linked to differences in individual behavior (Duda et al. 1999; Nussear et al. 2012; McGuire et al. 2014). Although previous studies provide insights towards potential factors that may affect burrow use, we lack a mechanistic understanding behind the role of these factors in driving heterogeneity in burrow use patterns in desert tortoises. A large impact of population stressors on refuge use can affect mating and foraging opportunities of desert tortoises and also reduce their likelihood of survival.

In this study, we combine data-sets from nine study sites in desert tortoise habitat, spanning more than 15 years to derive burrow use patterns of individuals in these populations. We first construct bipartite networks to infer their social associations due to asynchronous use of burrows. We then use generalized linear mixed models to explain mechanisms behind heterogeneity in burrow use behavior of individuals and effect of population stressors. As the desert tortoise is a long lived species, evaluating the impact of population stressors on burrow use patterns provides an efficient alternative to using traditional demographic metrics (such as mortality). We also investigate the use of burrows through a bipartite network model to identify why certain burrows are more popular than others in desert tortoise habitat. Overall, our analysis of refuge-based associations provide further insights into the structure and dynamics of social organization in a species traditionally considered as solitary and provides mechanisms behind individual variation within these associations.

## Methods

### Dataset

We combined datasets from nine study sites monitored from 1996 to 2014 across desert tortoise habitat in the Mojave desert of California, Nevada, and Utah (Fig. 1). Each site was monitored over multiple years, but not all sites were monitored in each year of the 15 year span. At each site, individuals were monitored at least weekly during their active season and at least monthly during winter months using radio telemetery. The total number of animals sampled and average number of observations per tortoise at each site is included in Supplementary Table S1. All tortoises were individually tagged, and during each tortoise encounter, data were collected to record the individual identifier, date of observation, GPS location, micro-habitat currently used by the animal (e.g., vegetation, pallet, or a burrow), any visible signs of injury or upper respiratory tract disease. As the dataset involved monitoring tagged individuals, it was not possible to record data blind. The unique burrow identification (id) was recorded for cases where an animal was located in a burrow. New burrow ids were assigned when an individual was encountered at a previously unmarked burrow.

**Fig. 1.**
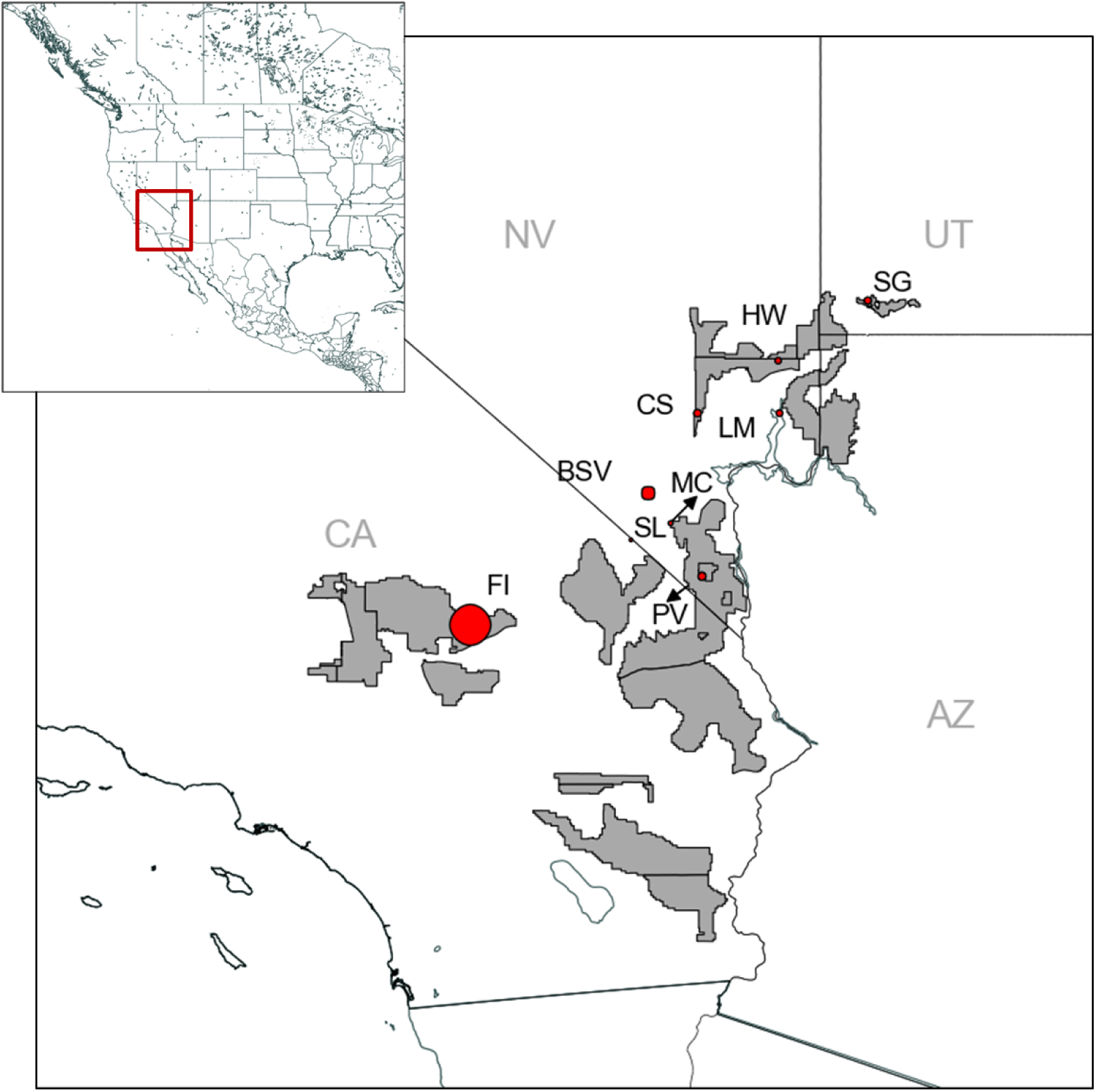
Critical habitat range of the desert tortoise within the Mojave desert, USA as determined by the US Fish and Wildlife Services in 2010 (http://www.fws.gov/). Critical habitat is defined as those geographical areas that contain physical or biological features essential to the conservation and management of the species (Department of the Interior 1973). Points represent centroids of survey sites where tortoises were monitored using radio-telemetry. Point size is proportional to the number of animals monitored at the site. Site abbreviations: BSV - Bird Spring Valley, CS - Coyote Springs, FI - Fort Irwin, HW - Halfway, LM - Lake Meade, MC - McCullough Pass, PV - Piute Valley, SG - St. George, SL - Stateline Pass

### Network Analysis

We constructed *bipartite networks* of asynchronous burrow use in desert tortoises for active (March - October) and inactive season (November - February) of each year at five sites (CS, HW, MC, PV, SL) where no translocations were carried out. An example of a burrow use bipartite network is shown in Fig. 2. The network consisted of burrow and tortoise nodes and undirected edges. An edge connecting a tortoise node to a burrow node indicated burrow use by the individual (Fig. 2). To reduce bias due to uneven sampling, we did not assign edge weights to the bipartite networks. Edges in a bipartite network always connect the two different node types, thus edges connecting two tortoise nodes or two burrow nodes are not permitted. Tortoise nodal degree in the bipartite network therefore denotes the number of unique burrows used by the individual and burrow nodal degree is the number of unique individuals visiting the burrow. Networks were generated using Networkx package in Python (Hagberg et al. 2008).

**Fig. 2.**
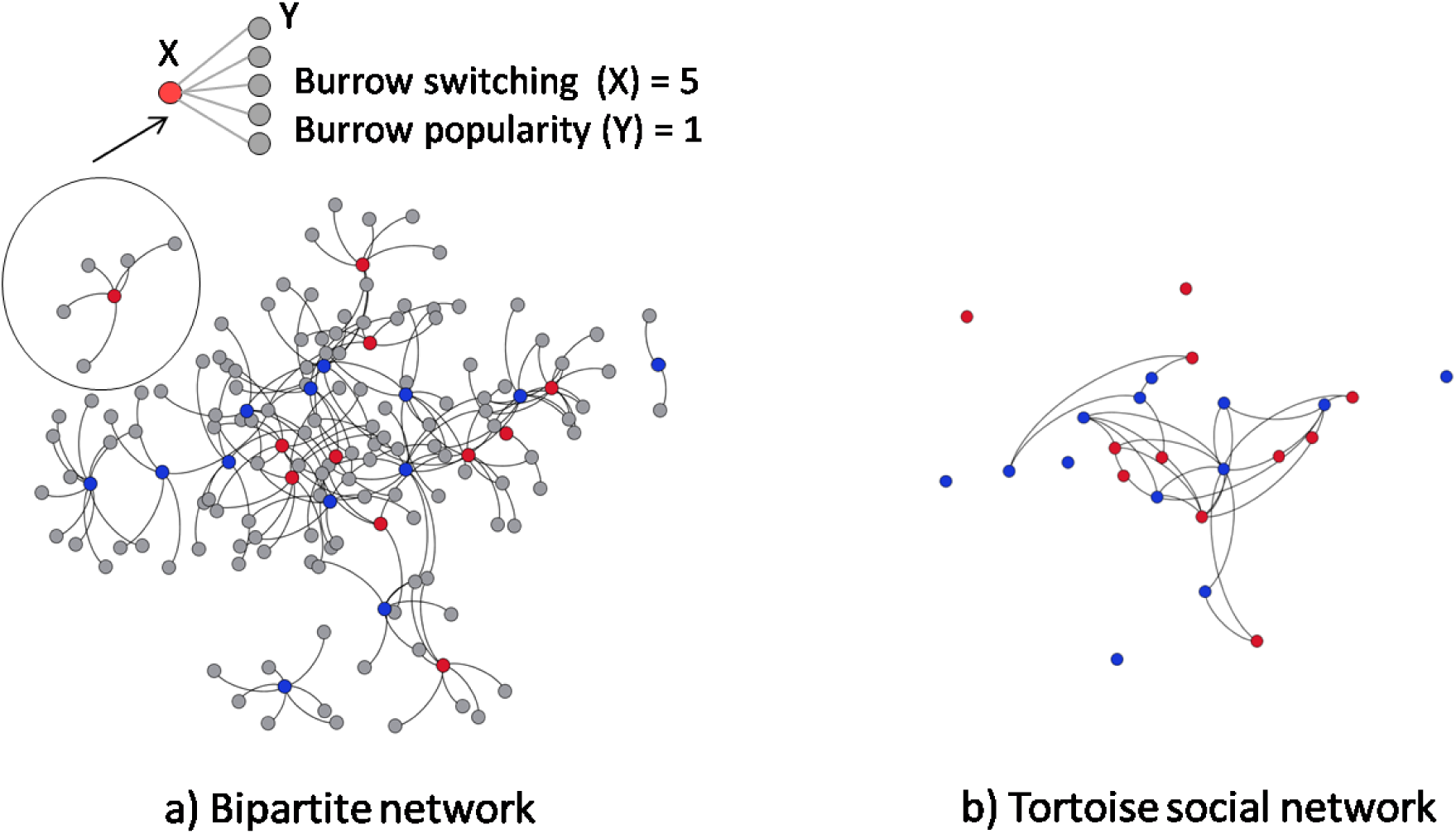
(a) Bipartite network of burrow use patterns at MC site during the year 2012. Node type indicated by color (Blue = adult males and red = adult females). Node positions are fixed using Yifan Hu’s multilevel layout in Gephi (Bastian et al. 2009). In this paper, we quantify burrow switching and burrow popularity as degree of tortoise nodes and burrow nodes, respectively, in the bipartite network. For example, burrow switching of the female tortoise X is five and burrow popularity of burrow Y is one. (b) Single-mode projection of the bipartite network into tortoise social network. Nodes with zero degree have been removed for clarity of illustration

We further examined the social structure of desert tortoises by converting the bipartite network into a single-mode projection of tortoise nodes (Tortoise social network, Fig. 2). For these tortoise social networks, we calculated network density, degree centralization, modularity, clustering coefficient, and homophily of individuals by degree and sex/age class. Network density is calculated as the ratio of observed edges to the total possible edges in a network (Scott and Carrington 2011). Degree centralization measures the variation in node degree across the network, such that high values indicate a higher heterogeneity in node degree and that a small proportion of nodes have a higher degree than the rest (Scott and Carrington 2011). Modularity measures the strength of the division of nodes into subgroups (Girvan and Newman 2002) and clustering coefficient measures the tendency of neighbours of a node to be connected (Bansal et al. 2009). The values of modularity and clustering coefficient can range from 0 to 1, and larger values indicate stronger modularity or clustering coefficient. We generated 1000 random network counterparts to each empirical network using double-edge swap operation in NetworkX (Hagberg et al. 2008) to determine if the observed network metrics were significantly different from random expectation. The generated random networks had the same degree sequence as empirical networks, but were random with respect to other network properties.

We next examined the spatial dependence of asynchronous burrow associations by using coordinates of burrows visited by tortoises to calculate centroid location of each tortoise during a particular season of a year. Distances between each tortoise pair (i, j) were then calculated as 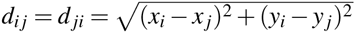 where (x, y) is the coordinate of tortoise centroid location. Pearson correlation coefficient was used to calculate the correlation between observed edges in social network and geographical distances between the tortoises. We compared the observed correlation to a null distribution of correlation values generated by randomly permuting spatial location of burrows 10,000 times and recalculating correlation between social associations and distance matrix for each permutation. Correlation were calculated using MantelTest package in Python (Carr 2015).

### Regression Analysis

We used generalized linear mixed regression models with Poisson distribution and log link function to assess burrow use patterns. To capture seasonal variation in burrow use, we aggregated the response counts over six periods (Jan-Feb, Mar-Apr, May-Jun, Jul-Aug, Sep-Oct and Nov-Dec). Patterns of burrow use were analyzed in two ways. First, we investigated factors affecting burrow switching, which we define as the number of unique burrows used by a tortoise in a particular sampling period. Second, we investigated burrow popularity, defined as the number of unique individuals using a burrow in a particular sampling period. Model variables used for each analysis are summarized in Table 1. All continuous model variables were centered (by subtracting their averages) and scaled to unit variances (by dividing by their standard deviation). This standard approach in multivariate regression modeling assigns each continuous predictor with the same prior importance in the analysis (Schielzeth 2010). All analyses were performed in R (version 3.0.2; R Development Core Team 2013).

#### Investigating burrow switching of desert tortoises

In this model, the response variable was burrow switching, defined as the total number of unique burrows used by desert tortoises during each sampling period. An individual was considered to be using a burrow if it was reported either inside a burrow or within 25 *m*^2^ grid around a burrow. The predictors included in the model are described in Table 1. In addition to the fixed effects, we considered three interactions in this model (i) sampling period × sex, (ii) sampling period × seasonal rainfall and (iii) local tortoise density × local burrow density. Tortoise identification and year × site were treated as random effects.

**Table 1.** Model variables considered to characterize burrow use patterns in the desert tortoise, *Gopherus agassizii*

| Variables | Variable type | Description |
| --- | --- | --- |
| **Tortoise attributes (Burrow switching model only)** |  |  |
| Sex/age class | Categorical | Three levels - adult males, adult females and non-reproductives (including neonates, juveniles and subadults) |
| Size | Continuous | Midline carapace length averaged over the year for each individual |
| **Burrow attributes (Burrow popularity model only)** |  |  |
| Burrow azimuth | Categorical | Direction in which burrow entrance faces forward. We converted the 1 to 360° range of possible azimuth values to eight categorical azimuth directions: Q1 (1-45), Q2 (46-90), Q3 (91-135), Q4 (136-180), Q5 (181-225), Q6 (226-270), Q7 (271-315) and Q8 (316-360) |
| Burrow surveyed age | Continuous | Number of years between the first report of burrow and current observation |
| Soil condition | Categorical | The soil conditions at the nine sites varied from sandy to mostly rocky. We therefore categorized burrow soil into four categories - mostly sandy, sand and rocky, mostly rocky and caliche and rocky |
| Percentage wash | Continuous | Percentage area covered by dry bed stream within 250 sqm area around burrow |
| Surface roughness | Continuous | See Inman et al. (2014) |
| Topographic position | Continuous | Index of landscape elevation around 250 sqm of burrow. High values are indicative of dry lakebeds or valley bottoms, and low values represent ridges and mountain tops. See Inman et al. (2014) for details. |
| **Environmental characteristics** |  |  |
| Sampling period | Categorical | The period of observation as described before. We divided a year into six periods of two months each |
| Seasonal rainfall* | Continuous | Total rainfall recorded at weather station nearest to the study site (in inches) during a particular sampling period |
| Temperature* | Continuous | Average, maximum and minimum temperature recorded at the weather station nearest to the study site and calculated over each sampling period in our model * |
| **Population stressors\*\*** |  |  |
| Tortoise health | Categorical | Burrow switching model only. Two categories - healthy and unhealthy |
| Residency status | Categorical | Burrow switching model only. Each individual was assigned one the five residency status for each sampling period - Control (C), Resident (R), Translocated (T), Ex-Resident (ER) or Ex-Translocated (ET) |
| Drought condition | Continuous | Average rainfall from November to February used as a proxy of drought condition for the following year |
| **Density condition** |  |  |
| Local tortoise density | Continuous | For burrow switching model: the average number of individuals found within 10,000 sqm grid around the focal tortoise each day of sampling period when the animal was surveyed. For burrow popularity model: number of individuals found in 10,000 sqm grid around the focal burrow averaged each surveyed day of the sampling period |
| Local burrow density | Continuous | For burrow switching model: the average number of active burrows in 10,000 sqm grid around the focal tortoise each day of the sampling period when the animal was reported. For burrow popularity model: the number of active burrows in 10,000 sqm grid around the focal burrow. A burrow was considered to be active if it was reported to be occupied at least once during the current or any previous sampling period |
| **Survey condition** |  |  |
| Sampling days | Continuous | Total survey days during the sampling period |
| Individual level bias | Continuous | Burrow switching model: Total number of days when the focal tortoise was reported using any burrow to account for any survey biases between individuals. Burrow popularity model: Total tortoises surveyed during the sampling period |

\*\* See text for details

#### Investigating burrow popularity

For this model, the response variable was burrow popularity defined as the total number of unique tortoises using a focal burrow in a sampling period. The predictors included in the model are also described in Table 1. In this model, we also tested for three interactions between predictors including (i) sampling period × seasonal rainfall, (ii) sampling period × local tortoise density, and (iii) local tortoise density × local burrow density. We treated burrow identification and year × site as random effects.

#### Population stressors

*Disease:* We considered tortoises exhibiting typical signs of URTD including nasal discharge, swollen (or irritated/ sunken) eyes, and occluded nares to be indicative of an unhealthy animal. As diagnostic testing was not the focus of the studies collecting the data, we were unable to confirm the infection status of individuals. Knowledge of confirmed infection status of animals, however, was not central to our study as our aim was to measure behavioral response of symptomatic individuals only. We included health condition in the regression model as a categorical variable with two levels - healthy and unhealthy. An individual was considered to be unhealthy if it was reported to display clinical signs of URTD at least once during the sampling period.

*Translocation:* We accounted for translocation in the regression model by giving each surveyed tortoise one of the following five residency status at each sampling period: Control (C), Resident (R), Translocated (T), Ex-resident (ER) or Ex-translocated (ET). Translocations were carried out at four (BSV, FI, LM, SG) out of nine sites in our dataset for purposes described in previous studies (Drake et al. 2012; Nussear et al. 2012). All animals native to the site were categorized as Controls (C) during sampling periods before translocation occurred. For sampling periods post translocation, all native animals were categorized as Residents (R), and introduced animals were categorized as Translocated (T). One year after translocation, translocated and resident tortoises were considered to be Ex-translocated (ET) and Ex-residents (ER), respectively, to account for potential acclimatization of introduced animals (Nussear et al. 2012). We note that one of the four translocation sites (SG) did not have native animals prior to translocation. No translocations were carried out at the rest of the five sites, so all animals surveyed at those sites were labeled as controls in all sampling periods.

*Drought:* The desert tortoise habitat in Mojave desert typically receives most of the rainfall during the winter season. We therefore used winter rainfall to assess drought conditions in desert tortoise habitat. We defined winter rain during a year as average rainfall from November to February and used it as a proxy of drought condition for the following year. We note that summer rainfall in desert tortoise habitat varies from west to east, where summer rainfall becomes a larger component of the total annual precipitation in East Mojave desert (Henen et al. 1998). Therefore, although we used winter rainfall as a proxy of drought conditions, we considered the effects of summer precipitation implicitly by including seasonal rainfall as a separate predictor (see Table1).

#### Model selection and validation

Following Harrell (2002) we avoided model selection to remove non-significant predictors and instead present results of our full model. Using the full model allows model predictions conditional on the values of all predictors and results in more accurate confidence interval of effects of interest (Harrell 2002). The Bayesian information criterion (BIC) of model selection was used to identify the best higher order interactions. A potential drawback of including all independent variables in the final model is multicollinearity. We therefore estimated Generalized Variance Inflation Factor (GVIF) values for each predictor. GVIF is a variant of traditional VIF used when any predictor in the model has more than 1 degree of freedom (Fox and Monette 1992). To make GVIF comparable across dimensions, Fox and Monette (1992) suggest using GVIF^(1/(2 Df))^ which we refer to as adjusted GVIF. We sequentially removed predictors with high adjusted GVIFs, recalculated adjusted GVIF, and repeated the process until all adjusted GVIF values in the model were below 3 (Zuur et al. 2010).

We carried out graphical diagnostics by inspecting the Pearson residuals for the conditional distribution to check if the models fit our data in each case. We detected under-dispersion in both the regression models. Under-dispersed models yield consistent estimates, but as equidispersion assumption is not true, the maximum-likelihood variance matrix overestimates the true variance matrix which leads to over-estimation of true standard errors (Winkelmann 2003). We therefore estimated 95% confidence intervals of fixed and random effects using bootstrapping procedures implemented in ‘bootMER’ function in package lme4.

We tested for the significance of fixed factors in both the models using likelihood ratio test (R function mixed from afex package (Singmann 2013)). For significant categorical predictors, we used Tukeys HSD (R function glht from the multcomp package, (Hothorn et al. 2008)) as a post-hoc test of significant pair-wise differences among means. All reported P-values of post-hoc tests are adjusted for multiple comparisons using the single-step method (Hothorn et al. 2008).

## Results

### Network Analysis

Bipartite networks of asynchronous burrow use across all sites demonstrated considerable variation in degree of tortoise nodes and burrow nodes (Fig. 3). Tortoises visited more unique burrows on average (4.03 ± 3.43 SD) and had a greater range of burrows visited in active seasons (1-9) than in inactive seasons (average = 1.46±0.72 SD, range = 1-5). Less than 40% of tortoises used more than one burrow during Nov-Feb (inactive) months (Fig. 3a). Most burrows in desert tortoise habitat were visited by a single tortoise during active and inactive season (Fig. 3b). Heterogeneity in the number of animals visiting burrows, however, tended to be slightly more during the months of March-November than November-February (active = 1.21±0.56 SD, inactive = 1.08±0.35 SD).

**Fig. 3.**
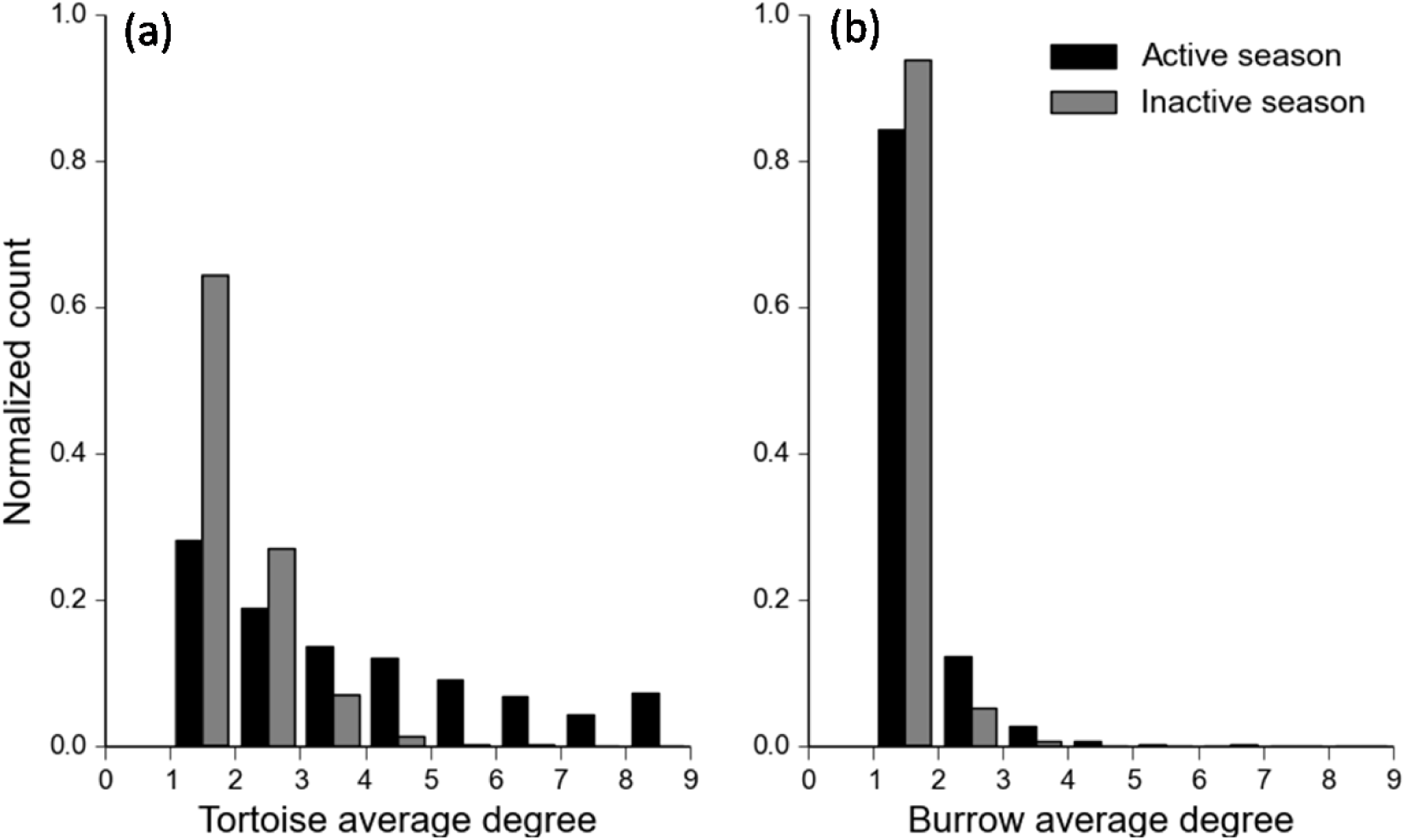
Frequency distribution of (a) Tortoise degree i.e., unique burrows used by desert tortoises and (b) Burrow degree i. e., unique tortoises visiting burrows during active (Mar-Oct) and inactive (Nov-Feb) seasons, excluding zero degree nodes. Values are averaged over each surveyed year and study site. y-axis represents normalized frequency counts of tortoises/burrows

The tortoise social network (constructed as a single mode projection of tortoise nodes from the bipartite network) demonstrated moderate clustering coefficient (0.36 ± 0.21 SD) and modularity (0.53 ± 0.15 SD). Twenty three of the 24 social networks we analyzed had higher clustering coefficient and 18 social networks were more modular than random networks (Supplementary Table S3). Thirteen social networks out of the total 24 demonstrated significant degree homophily (when nodes with similar degree tend to be connected) and 11 of those had positive associations (Supplementary Table S3). Positive degree homophily suggests that tortoises using many unique burrows often use the same set of burrows and are therefore connected in the social network. Tortoise social networks also had a moderate positive degree centralization which indicates a small subset of individuals used more burrows than the rest in the sampled population. Within sexes, positive degree centralization was observed both within males (0.20 ± 0.08 SD) and females (0.17 ± 0.06 SD). Homophilic association by sex ranged from -0.6 to 0.11 indicating a preference for one sex to associate with the opposite. These negative sexwise associations, however, were not different than those expected by chance.

The association between tortoises in their social network was inversely correlated with geographical distances between them, indicating that individuals closer to each other preferred using the same set of burrows. The magnitude of correlation ranged from -0.22 to -0.89 with an average value of -0.49 (Fig. 4). The P-value of the permutation test for all sites across active seasons of all surveyed years was less than 0.05, indicating a significant effect of geographical location on social associations (Supplementary Table S4). This result of spatial constraints driving social interactions is not surprising as geographical span of surveyed sites were much larger (> 1500m) than the normal movement range of desert tortoises (Franks et al. 2011). However, the moderate value of correlations suggest other factors (such as environmental, social, density) could play an important role in desert tortoise’s asynchronous burrow associations.

**Fig. 4.**
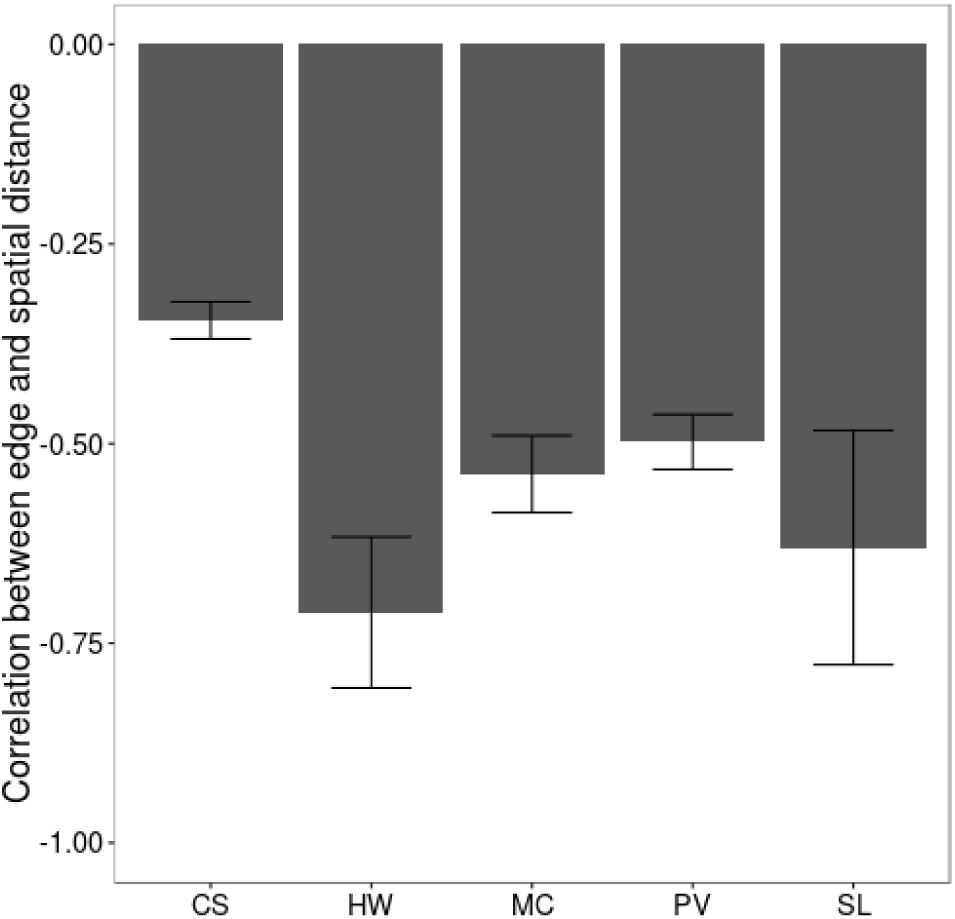
Spatial constraints on asynchronous burrow associations during active seasons at study sites with control animals. At each site, correlation is calculated between geographical distance and edge occurrence in tortoise social network, and averaged over each surveyed year. Error bars represent standard errors with n=8 (CS), n=3 (HW), n=2 (MC), n=7 (PV) and n=2 (SL). P-value associated with each correlation measure is < 0.05

### Regression Analysis

Based on the observed heterogeneity in bipartite networks, we next investigated the relative effect of natural variables and population stressors on burrow switching patterns of desert tortoises (*viz* degree of animal nodes in bipartite networks) and popularity of burrows in desert tortoise habitat (*viz* degree of burrow nodes in bipartite networks). Supplementary Table S5 presents the best models of BIC values for interactive predictors that explain burrow switching in desert tortoises and burrow popularity. The three interactions tested for burrow switching models were sampling period × sex, sampling period × seasonal rainfall and local tortoise density × local burrow density. We tested all possible combinations of the three interactions. The best model contained an interaction of sampling period × seasonal rainfall (Supplementary Table S5). For the burrow popularity model, we tested all possible combinations of the sampling period × seasonal rainfall, the sampling period × the local tortoise density and the local tortoise density × local burrow density interactions. The best model included the sampling period × the local tortoise density and the local tortoise density × the local burrow density interaction term.

Multicollinearity tests revealed all three measures of temperature (average, max and min) to have adjusted GVIF values of >3. The three predictors were therefore dropped from both the models. We also removed the sampling period × tortoise density interaction from the burrow popularity model as it inflated adj GVIF value of tortoise density to >3. σ^2^ estimate of tortoise identification and burrow identification random effect was negligible (tortoise identification: σ^2^ = 0, CI = 0-0.004, burrow identification: σ^2^ = 0, CI = 0-0.01). Both random effects were therefore removed from the regression models.

### Effect of animal attributes

Sex/age class had a significant effect on burrow switching 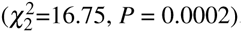. Overall, adults used more unique burrows than non-reproductives. Among adults, males used a slightly higher number of unique burrows than females (Fig. 5). There was no effect of body size on individuals’ burrow switching behavior 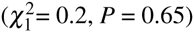.

**Fig. 5.**
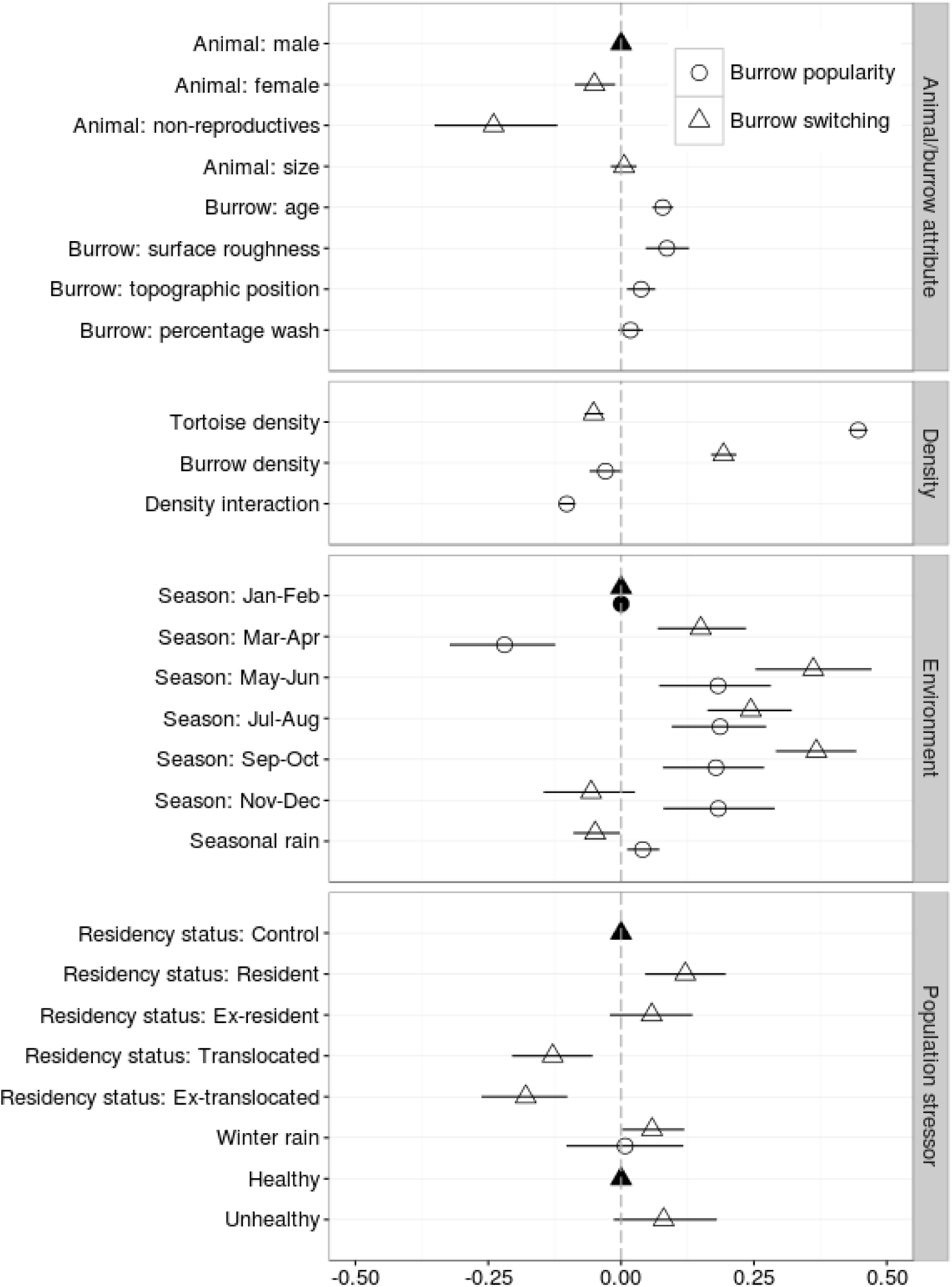
The effect of various predictors on the two models of burrow use patterns in desert tortoises. Error bars indicate 95% confidence intervals around the estimated coefficient value. For continuous predictors, the vertical dashed line indicates no effect - positive coefficients indicate increase in burrow popularity/switching with increase in predictor value; negative coefficients indicate decrease in burrow popularity/switching with higher values of predictors. For each categorical predictor, the base factor (solid data points) straddles the vertical line at 0 and appears without a 95% CI. Positive and negative coefficients for categorical predictors denote increase and decrease, respectively, in burrow popularity/switching relative to the base factor

### Effect of burrow attributes

Out of the six burrow attributes included in the model, burrow age and surface roughness around burrow had the highest impact on burrow popularity, i.e., number of unique individuals visiting the burrow (burrow age: 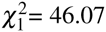, *P* < 0.0001, surface roughness: 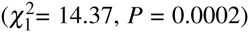. Burrow popularity was positively correlated with surface roughness indicating that burrows in flat sandy areas were visited by fewer unique tortoises than burrows in rough rocky areas (Fig. 5). Older burrows were visited by more unique individuals, with burrow popularity increasing *e*^0.08^ times with each increment of age (Fig. 5). Burrows in areas with higher topographical position as indicated by GIS raster images were also more popular 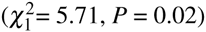.

### Effect of environmental conditions

Sampling period had a large effect on number of unique burrows used by desert tortoises 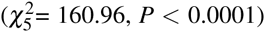 as well as on burrow popularity 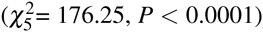 as compared to other model predictors. Burrow switching of desert tortoises was highest during the months of May-June and September-October when they are typically more active, and lowest in winter months (Fig. 5). In the late summer (July-August), tortoises demonstrated slightly lower burrow switching than during the active season, but higher than the winter season. Within a particular year, the direction of the effect of seasonal rainfall varied across different sampling periods (sampling period × seasonal rain: 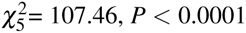). For example, high rainfall during the months of March-April reduced burrow switching in desert tortoises. On the other hand, individuals exhibited higher burrow switching with higher rain during the months of July-August (Supplementary Fig. S3b).

In contrast to the large variation in individuals' burrow switching behavior between sampling periods, popularity of burrows did not vary during a large portion of the year (May - December). Total unique animals visiting burrows tended to be lower in the months of January-February and March-April, as compared to other months of the year (Fig. 5, S4c). Seasonal rainfall had a positive correlation with burrow popularity 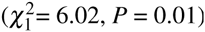.

### Effect of density conditions

An increase in the number of active burrows around individuals promoted burrow switching, whereas an individual used fewer burrows when there were more tortoises in the vicinity (Fig. 5). In the burrow popularity model, higher tortoise density around burrows increased number of individuals visiting these burrows (Fig. 5). There was a significant interactive effect of the two density conditions on burrow popularity 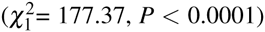 – increase in burrow popularity with higher tortoise density was lower when there were more burrows in the vicinity of the focal burrow (Supplementary Fig. S4d).

### Effect of population stressors

Population stressors of drought, health and translocation had variable influences on burrow switching of desert tortoises (Fig. 5, Supplementary Fig. 5). As compared to residents and controls, translocated animals demonstrated lower burrow switching during the year of translocation and also in the subsequent years (Fig. 5, Supplementary Fig. S5a). We did not find any differences between burrow switching levels of individuals exhibiting clinical signs of URTD and clinically healthy individuals 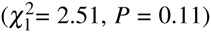. Burrow switching levels of all surveyed animals during drought years (indicated by lower winter rainfall), however, tended to be slightly lower in comparison to non-drought years (burrow switching: 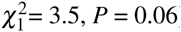).

## Discussion

Although direct social interactions among solitary species are relatively infrequent, individual preference for certain shared refuge and foraging spaces may lead to a highly structured social system (Leu et al. 2011). In such species, knowledge of social network structure formed through refuge or forage associations can identify key influential individuals (Fortuna et al. 2009; Leu et al. 2011), and provide early-warning signals for environmental (or anthropogenic) disturbances (Jachowski et al. 2012; Moule et al. 2015) that may ultimately affect population fitness. In this study, we infer social associations between individuals of a relatively solitary species, the desert tortoise, by their asynchronous use of burrows. While descriptive approaches are common in the field of animal social networks (Pinter-Wollman et al. 2013), we sought to gain a mechanistic understanding behind individual variation in burrow-use associations of desert tortoises. The degree of an individual in a bipartite network has biological and ecological importance as it indicates a decision to switch refuges. Refuge switching is associated with a tradeoff between the costs of increasing exposure to heat, predators, increased risk of infection, and the benefits of finding food and mates. The outcome of observed refuge switching patterns is important as theoretical models predict reduced survival of populations due to suboptimal refuge use decisions (Cooper 2015). Modeling optimal burrow switching that maximizes fitness in desert tortoises is challenging as it is difficult to quantify fitness costs in a long-lived species. Our study instead provides an approach to build baseline models of burrow use patterns. Any large deviation to baseline levels may indicate lower survival, foraging, and reproductive success for tortoises and thus burrow switching can serve as an immediate indicator of population stressors affecting long-term fitness consequences.

We show that social networks in desert tortoises formed due to burrow use preferences cannot be explained by random associations. In several wildlife systems, spatial constraints can play a large role in shaping social networks (Davis et al. 2015), and non-random associations may not be definitive evidence of social organization in a population. Desert tortoise social associations, however, were only moderately correlated to spatial distances, which corroborates earlier studies that report social organization in desert tortoises (Niblick et al. 1994; Bulova 1997). In general, the social networks were also clustered (0.23-0.59) and modular (0.34 - 0.68). However, higher clustering coefficient values have been reported in other social species [e.g, 0.54-0.57 in bottlenose dolphins (Mann et al. 2012), 0.57 - 0.87 in guppies (Croft et al. 2004), 0.81 in squirrels (Manno 2008), 0.57-0.67 in primates (Pasquaretta et al. 2014)] and even in a few relatively solitary species that have been studied [e.g., 0.7 in raccoons (Hirsch et al. 2013), 0.59 in brushtail possum (Porphyre et al. 2011)]. The low (but significant) clustering coefficient value in desert tortoise social networks suggests that they do not form tight social bonds as compared to other social wildlife species. In social species, the network structure is known to affect population stability (Kurvers et al. 2014) and resistance to disease outbreaks (Cross et al. 2004; Godfrey et al. 2009; MacIntosh et al. 2012). Modular social networks of desert tortoises in particular can have important implications in the spread and persistence of infections. For example, few connections between communities in a social network can effectively localize new infections to a few individuals. For chronic infections such as URTD, these pockets of infection, however, can serve as sources of re-infection to other uninfected communities, eventually leading to persistent infection across the entire population.

Our analysis of burrow use heterogeneity in desert tortoises reveals that the period of the year and density of burrows around an individual are the main drivers behind the individual’s burrow switching decision. Low burrow switching levels in tortoises during winter and summer months reflects reduced movement to avoid severe weather conditions (Eubanks et al. 2003). Individuals visit more burrows in the months of May-June and September-October which coincides with high activity of nesting and mating in adults. Seasonal rainfall also influences burrow switching in desert tortoises. Tortoises use fewer burrows in high rainfall conditions in March-April months, which possibly reflects reduced activity during cold weather associated with spring storms. Infrequent summer rains, on the other hand, increase tortoise activity as individuals emerge from burrows to rehydrate (Nagy and Medica 1986; Peterson 1996). Our results of high burrow switching during summer rains (July-August) are consistent with these reports of increased activity. We also find that non-reproductive tortoises use fewer burrows than adults, which may reflect differences in costs and benefits associated with switching burrows. Leaving a refuge can present a greater risk to non-reproductives that are more vulnerable to predation (Wilson 1991), are prone to thermal stress due to their smaller size (Mushinsky et al. 2003), and do not benefit from the mating opportunities gained by burrow switching. Indeed, previous studies have found juveniles forage closer to their burrows and minimize time spent out of burrows (Mcrae et al. 1981; Mushinsky et al. 2003; Halstead et al. 2007). Future studies and management plans of desert tortoises may consider differences in burrow switching between different non-reproductive tortoises in order to mitigate increased predation risk by pervasive predators such as ravens.

While it has been shown that a small fraction of burrows in desert tortoises are visited by multiple animals (Bulova 1994; Harless et al. 2009), the mechanisms behind burrow popularity were previously unknown. Our results suggest that popular burrows can be identified using certain burrow characteristics such as surrounding topographical variables and age. As true burrow age is often hard to determine, we demonstrate the use of historical survey data to estimate proxy age of burrows. Once identified, these popular burrows can be surveyed throughout the year as there is only a minor effect of time of the year and seasonal rainfall on burrow popularity. Knowledge of active and popular refuges can have two important implications for the conservation and management of wildlife species. First, population density estimates usually rely on observations of animals located outside refuge space (Witmer 2005). For species that spend most of the time in a year in a refuge, survey of popular refuges can augment the current survey methods to get a more accurate estimate of population density. Secondly, declines of popular refuges can indicate reduced social interactions and mating opportunities for individuals. Reduced refuge popularity can also be indicative of higher mortality risk - Esque et al. (2010) found higher mortality of desert tortoise in flat open areas where burrows, as our results indicate, are less popular compared to rough higher elevation sites. Active popular burrows can therefore be used (a) as sentinels of population health and (b) to identify critical core habitat for conservation and adaptive management of a wildlife species.

Of three potential population stressors that we included in our model (disease, drought, translocation), translocation caused a change in burrow switching behavior of desert tortoises. Although translocated animals are known to have high dispersal tendencies (Nussear et al. 2012; Hinderle et al. 2015) and hence are expected to encounter and use more burrows, we found translocated individuals use fewer unique burrows than residents. Our results are supported by evidence of translocated tortoises spending more time on the surface and taking shelter under vegetation rather than using burrows (Hinderle 2011). The use of fewer burrows coupled with high dispersal rates can increase exposure of translocated animals to thermal stress and dehydration, potentially increasing mortality. Therefore, to improve translocation success, a fruitful area of investigation for future research will be to determine potential causes of this change in burrow use behavior in translocated tortoises. We used winter rain as a proxy of drought conditions as the Western Mojave receives most of its annual rainfall during the months of November-February and is important for the availability of food for desert tortoises in the spring (Duda et al. 1999; Lovich et al. 2014). Our results show a slight (but not significant) reduction in burrow use by tortoises during drought years. Reduced burrow switching may correspond to smaller home-ranges of desert tortoises observed during drought years (Duda et al. 1999). Low winter rainfall condition is also known to increase predation of desert tortoises due to diminished prey resources (Peterson 1994; Esque et al. 2010). Lower burrow use during drought years can be therefore a behavioral response of desert tortoises to avoid predation or to reduce energy expenditure and water loss in years of low resource availability (Nagy and Medica 1986). Contrary to previous studies (McGuire et al. 2014), we did not find any effect of disease on burrow use behavior, possibly because we could not distinguish severe clinical signs with milder forms in our data. Although there was no evidence of disease influencing burrow use behavior in the present study, we note that it is likely for burrow use behavior (and in particular the burrows themselves) to drive infectious disease patterns in desert tortoises either directly, through cohabitation instances, or indirectly, by serving as focal sites of social interactions.

## Conclusions

Our study demonstrates non-random associations in desert tortoises based on refuge use patterns. We formulate statistical models of burrow switching and popularity of burrows to investigate the mechanisms including environmental, topographical, density factors, and population stressors behind refuge use preferences of desert tortoises. In combination, these models help infer the mechanisms behind heterogeneity in refuge use from the perspective of individuals as well as from the perspective of the refuges. This approach is particularly useful for species that are not overtly gregarious. For these species, refuge switching often correlates to reproductive and foraging success, and patterns of refuge use can be an important aspect to consider before implementing any management or conservation strategy. For example, popular refuges can be used to identify core habitat areas. In addition, sudden changes in the refuge switching behavior of individuals can be used as an early warning signal of disturbances that may ultimately affect population fitness. More broadly, our study provides insights towards the presence of and mechanisms behind non-random social structure and individual variation in a relatively solitary species by analyzing refuge-based associations. The structure of networks in social species is known to affect population stability and resilience to infectious diseases. Future studies are needed to establish such functional roles of social networks in relatively solitary species.

## Acknowledgments

We thank Phil Medica for feedback on a previous version of this manuscript. We thank Ian T. Carroll and anonymous reviewers for helpful comments to improve the quality of the paper. We thank Clarence Everly, Department of Defense, Ft. Irwin National Training Center for the support in acquiring the data analyzed in this paper. Any use of trade, product, or firm names is for descriptive purposes only and does not imply endorsement by the U.S.Government.

## Ethical approval

All applicable international, national, and/or institutional guidelines for the care and use of animals were followed. This article does not contain any studies with human participants performed by any of the authors.

## References

Bansal S, Khandelwal S, Meyers LA (2009) Exploring biological network structure with clustered random networks. BMC Bioinformatics 10:405

Bastian M, Heymann S, Jacomy M (2009) Gephi: An open source software for exploring and manipulating networks, https://gephi.org

Behringer DC, Butler IVMJ (2010) Disease avoidance influences shelter use and predation in Caribbean spiny lobster. Behav Ecol Sociobiol 64:747–755

Berger-Tal O, Polak T, Oron A, Lubin Y, Kotler BP, Saltz D (2011) Integrating animal behavior and conservation biology: a conceptual framework. Behav Ecol 22:236–239

Brown MB, Schumacher IM, Klein PA, Harris K, Correll T, Jacobson ER (1994) *Mycoplasma agassizii* causes upper respiratory tract disease in the Desert Tortoise. Infect Immun 62:4580–4586

Bulova SJ (1994) Patterns of burrow use by desert tortoises: gender differences and seasonal trends. Herpetol Monogr 8:133–143

Bulova SJ (1997) Conspecific chemical cues influence burrow choice by desert tortoises (*Gopherus agassizii*). Copeia 1997:802–810

Carr JW (2015) MantelTest, http://jwcarr.github.io/MantelTest

Carter GG, Wilkinson GS (2013) Food sharing in vampire bats: reciprocal help predicts donations more than relatedness or harassment. Proc R Soc B 280:2012–2573

Cooper WE Jr (2015) Escaping from predators: an integrative view of escape decisions. Cambridge University Press, Cambridge

Croft DP, Krause J, James R (2004) Social networks in the guppy (*Poecilia reticulata*). Proc R Soc Lond B 271:S516–S519

Cross PC, Lloyd-smith JO, Bowers JA, Hay CT, Hofmeyr M, Getz WM (2004) Integrating association data and disease dynamics in a social ungulate: bovine tuberculosis in African buffalo in the Kruger National Park. Ann Zool Fenn 41:879–892

Davis S, Abbasi B, Shah S, Telfer S, Begon M (2015) Spatial analyses of wildlife contact networks. J R Soc Interface 12: 2014–1004

Department of the Interior (1973) Endangered species act. Technical report, U.S. Fish and Wildlife Service, Washington, DC

Department of the Interior (2011) Revised recovery plan for the Mojave population of the desert tortoise (*Gopherus agassizii*). Technical report, U.S. Fish and Wildlife Service, Sacramento

Dodd KC, Seigel RA (1991) Relocation, repatriation, and translocation of amphibians and reptiles: are they conservation strategies that work? Herpetologica 47:336–350

Drake KK, Nussear KE, Esque TC, Barber AM, Vittum KM, Medica PA, Tracy CR, Hunter KW (2012) Does translocation influence physiological stress in the desert tortoise? Anim Conserv 15:560–570

Duda JJ, Krzysik AJ, Freilich JE (1999) Effects of drought on desert tortoise movement and activity. J Wildlife Manage 63:1181–1192

Esque TC, Nussear KE, Drake KK et al (2010) Effects of subsidized predators, resource variability, and human population density on desert tortoise populations in the Mojave Desert, USA. Endanger Species Res 12:167–177

Eubanks JO, Michener WK, Guyer C (2003) Patterns of movement and burrow use in a population of gopher tortoises (*Gopheruspolyphemus*). Herpetologica 59:311–321

Fortuna MA, Popa-Lisseanu AG, Ibáñfiez C, Bascompte J (2009) The roosting spatial network of a bird-predator bat. Ecology 90:934–944

Fox J, Monette G (1992) Generalized collinearity diagnostics. J Am Stat Assoc 87:178–183

Franks BR, Avery HW, Spotila JR (2011) Home range and movement of desert tortoises Gopherus agassizii in the Mojave Desert of California, USA. Endanger Species Res 13:191–201

Germano JM, Bishop PJ (2009) Suitability of amphibians and reptiles for translocation. Conserv Biol 23:7–15

Girvan M, Newman MEJ (2002) Community structure in social and biological networks. P Natl Acad Sci USA 99:7821–7826

Godfrey SS (2013) Networks and the ecology of parasite transmission: A framework for wildlife parasitology. Int J Parasitol Parasites Wildl 2:235–245

Godfrey SS, Bull CM, James R, Murray K (2009) Network structure and parasite transmission in a group living lizard, the gidgee skink, Egernia stokesii. Behav Ecol Sociobiol 63:1045–1056

Gough HM, Gascho Landis AM, Stoeckel JA (2012) Behaviour and physiology are linked in the responses of freshwater mussels to drought. Freshwater Biol 57:2356–2366

Griffin RH, Nunn CL (2011) Community structure and the spread of infectious disease in primate social networks. Evol Ecol 26:779–800

Hagberg AA, Schult DA, Swart PJ (2008) Exploring network structure, dynamics, and function using NetworkX. Proceedings of the 7th Python in Science Conference (SciPy2008) 836:11–15

Halstead BJ, McCoy ED, Stilson TA, Mushinsky HR (2007) Alternative foraging tactics of juvenile gopher tortoises (*Gopherus Polyphemus*) examined using correlated random walk models. Herpetologica 63:472–481

Harless ML, Walde AD, Delaney DK, Pater LL, Hayes WK (2009) Home range, spatial overlap, and burrow use of the desert tortoise in the West Mojave Desert. Copeia 2009:378–389

Harrell FE (2002) Regression modeling strategies: with applications to linear models, logistic and ordinal regression, and survival analysis, 1st edn. Springer, London

Henen BT, Peterson CC, Wallis IR, Berry KH, Nagy KA (1998) Effects of climatic variation on field metabolism and water relations of desert tortoises. Oecologia 117:365–373

Hinderle D (2011) Desert tortoises (*Gopherus agassizii*) and translocation: homing, behavior, habitat and shell temperature experiments. Technical report, San Diego State University, San Diego

Hinderle D, Lewison RL, Walde AD, Deutschman D, Boarman WI (2015) The effects of homing and movement behaviors on translocation: Desert tortoises in the western Mojave Desert. J Wildlife Manage 79:137–147

Hirsch BT, Prange S, Hauver SA, Gehrt SD (2013) Raccoon social networks and the potential for disease transmission. PLoS ONE 8:e75830

Hothorn T, Bretz F, Westfall P (2008) Simultaneous inference in general parametric models. Biometrical J 50:346–363

Inman RD, Nussear KE, Esque TC, Vandergast AG, Hathaway SA, Wood DA, Barr KR, Fisher RN (2014) Mapping habitat for multiple species in the Desert Southwest. US Geological Survey, Reston

Jachowski DS, Slotow R, Millspaugh JJ (2012) Physiological stress and refuge behavior by african elephants. PLoS ONE 7:e31818

Jacobson ER, Brown MB, Wendland LD, Brown DR, Klein PA, Christopher MM, Berry KH (2014) Mycoplasmosis and upper respiratory tract disease of tortoises: a review and update. Vet J 201:257–64

Kerr GD, Bull CM (2006) Movement patterns in the monogamous sleepy lizard (*Tiliqua rugosa*). Effects of gender, drought, time of year and time of day. J Zool 269:137–147

Kurvers RHJM, Krause J, Croft DP, Wilson ADM, Wolf M (2014) The evolutionary and ecological consequences of animal social networks: emerging issues. Trends Ecol Evol 29:326–35

Leu ST, Kappeler PM, Bull CM (2010) Refuge sharing network predicts ectoparasite load in a lizard. Behav Ecol Sociobiol 64:1495–1503

Leu ST, Kappeler PM, Bull CM (2011) The influence of refuge sharing on social behaviour in the lizard *Tiliqua rugosa*. Behav Ecol Sociobiol 65:837–847

Longshore KM, Jaeger JR, Sappington JM (2003) Desert tortoise (*Gopherus agassizii*) survival at two eastern Mojave Desert sites: death by short-term drought? J Herpetol 37:169–177

Lovich JE, Yackulic CB, Freilich J, Agha M, Austin M, Meyer KP, Arundel TR, Hansen J, Vamstad MS, Root SA (2014) Climatic variation and tortoise survival: Has a desert species met its match? Biol Conserv 169:214–224

Lusseau D, Wilson B, Hammond PS, Grellier K, Durban JW, Parsons KM, Barton TR, Thompson PM (2006) Quantifying the influence of sociality on population structure in bottlenose dolphins. J Anim Ecol 75:14–24

MacIntosh AJJ, Jacobs A, Garcia C, Shimizu K, Mouri K, Huffman MA, Hernandez AD (2012) Monkeys in the middle: parasite transmission through the social network of a wild primate. PloS ONE 7:e51144

Mann J, Stanton MA, Patterson EM, Bienenstock EJ, Singh LO (2012) Social networks reveal cultural behaviour in tool-using using dolphins. Nat Commun 3:980

Manno TG (2008) Social networking in the Columbian ground squirrel, *Spermophilus columbianus*. Anim Behav 75:1221–1228

McGuire JL, Smith LL, Guyer C, Yabsley MJ (2014) Effects of mycoplasmal upper-respiratory-tract disease on movement and thermoregulatory behavior of gopher tortoises (*Gopherus Polyphemus*) in Georgia, USA. J Wildlife Dis 50:745–756

Mcrae WA, Landers JL, Garner JA (1981) Movement patterns and home Range of the gopher tortoise. Am Midl Nat 106:165–179

Morris DW, Kotler BP, Brown JS, Sundararaj V, Ale SB (2009) Behavioral indicators for conserving mammal diversity. Ann NY Acad Sci 1162:334–56

Moule H, Michelangeli M, Thompson MB, Chapple DG (2015) The influence of urbanization on the behaviour of an Australian lizard and the presence of an activity-exploratory behavioural syndrome. J Zool 298:103–111

Mushinsky HR, Stilson TA, McCoy ED (2003) Diet and dietary preference of the juvenile gopher tortoise (*Gopherus Polyphemus*). Herpetologica 59:475–483

Nagy KA, Medica PA (1986) Physiological ecology of desert tortoises in southern Nevada. Herpetologica 42:73–92

Niblick HA, Rostal DC, Classen T (1994) Role of male-male interactions and female choice in the mating system of the desert tortoise, *Gopherus agassizii*. Herpetol Monogr 8:124–132

Nussear KE, Tracy CR, Medica PA, Wilson DS, Marlow RW, Corn PS (2012) Translocation as a conservation tool for Agassiz’s desert tortoises: survivorship, reproduction, and movements. J Wildlife Manage 76:1341–1353

Pasquaretta C, Leve M, Claidiere N et al (2014) Social networks in primates: smart and tolerant species have more efficient networks. Sci Rep 4:7600

Peterson CC (1994) Different rates and causes of high mortality in two populations of the threatened desert tortoise *Gopherus agassizii*. Biol Conserv 70:101–108

Peterson CC (1996) Anhomeostasis: seasonal water and solute in two relations of the desert populations tortoise (*Gopherus agassizil*) during Chronic Drought. Physiol Zool 69:1324–1358

Pinter-Wollman N, Hobson EA, Smith JE et al (2013) The dynamics of animal social networks: analytical, conceptual, and theoretical advances. Behav Ecol 25:242–255

Porphyre T, McKenzie J, Stevenson MA (2011) Contact patterns as a risk factor for bovine tuberculosis infection in a free-living adult brushtail possum *Trichosurus vulpecula* population. Prev Vet Med 100:221–230

Sandmeier FC, Tracy CR, DuPré S, Hunter K (2009) Upper respiratory tract disease (URTD) as a threat to desert tortoise populations: A reevaluation. Biol Conserv 142:1255–1268

Schielzeth H (2010) Simple means to improve the interpretability of regression coefficients. Method Ecol Evol 1:103–113

Scott J, Carrington P (eds) (2011) The SAGE Handbook of Social Network Analysis. SAGE publications, London

Singmann H (2013) afex: analysis of factorial experiments, https://cran.r-project.org/web/packages/afex

van Gils JA, Kraan C, Dekinga A, Koolhaas A, Drent J, de Goeij P, Piersma T (2009) Reversed optimality and predictive ecology: burrowing depth forecasts population change in a bivalve. Biol Lett 5:5–8

Vander Wal E, Paquet PC, Andres JA (2012) Influence of landscape and social interactions on transmission of disease in a social cervid. Mol Ecol 21:1271–82

Wilson DS (1991) Estimates of survival for juvenile gopher tortoises, *Gopherus polyphemus*. J Herpetol 25:376–379

Wilson DS, Morafka DJ, Tracy CR, Nagy KA (1999) Winter activity of juvenile desert tortoises (*Gopherus agassizii*) in the Mojave Desert. J Herpetol 33:496–501

Winkelmann R (2003) Econometric analysis of count data. Springer Science & Business Media, Heidelberg

Witmer GW (2005) Wildlife population monitoring: some practical considerations. Wildlife Res 32:259–263

Zuur A, Ieno E, Elphick CS (2010) A protocol for data exploration to avoid common statistical problems. Method Ecol Evol 1:3–14

